# The complex geography of domestication of the African rice *Oryza glaberrima*

**DOI:** 10.1101/321851

**Authors:** Jae Young Choi, Katherine Dorph, Michael D. Purugganan

## Abstract

While the domestication history of Asian rice has been extensively studied, details of the evolution of African rice remains elusive. The inner Niger delta has been suggested as the center of origin but molecular data for its support are lacking. Here, we present the most comprehensive analysis to date on the evolutionary and domestication history of African rice. By analyzing whole genome re-sequencing data from 282 individuals in domesticated African rice *Oryza glaberrima* and its progenitor *O. barthii*, we hypothesize a non-centric domestication origin for African rice. Our analyses show geographically based population structure in *O. glaberrima*, as well as significant evidence of admixture between genetic groups. Furthermore, we have evidence that the previously hypothesized *O. barthii* progenitor populations in West Africa have evolutionary signatures similar to domesticated rice and carried causal domestication mutations, suggesting those progenitors may actually represent feral wild-domesticated hybrid rice. Demography modeling suggested the inland *O. glaberrima* had a protracted period of bottlenecking that preceded the coastal population by 800–1,800 years. Phylogeographic analysis of genes involved in the core domestication process suggests that the origins of causal domestication mutations could be traced to wild progenitors in multiple different locations in West and Central Africa. Based on our evidence, we hypothesize *O. glaberrima* was not domesticated from a single centric location but was rather a diffuse process where multiple regions contributed key alleles for different domestication traits.

**Author Summary:** For many crops it is not clear how they got domesticated from their wild progenitors. Transition from a wild to domesticated state required a series of genetic changes, and studying the evolutionary origin of these domestication-causing mutations are key to understanding the domestication origins of a crop. Moreover, population relationships within a crop holds insight into the evolutionary history of domestication and whether there was gene flow between different genetic groups. In this study, we investigate the domestication history of *Oryza glaberrima*, a rice species that was domesticated in West Africa independently from the Asian rice species *O. sativa*. Using genome-wide data from a large sample of domesticated and wild African rice samples we did not find evidence that supported the established domestication model for *O. glaberrima*—a single domestication origin. Rather, our evidence suggests the domestication process for African rice was initiated in multiple regions of West Africa, caused potentially by the local environmental and cultivation preference of people. Admixture between different genetic groups had facilitated the exchange and spread of core domestication mutations. Hence domestication of African rice was a multi-regional process.

## Introduction

Domestication of crop species represents a key co-evolutionary transition, in which wild plant species were cultivated by humans and eventually gave rise to new species whose propagation was dependent on human action [1–3]. The evolutionary origin(s) of various crop species have been the subject of considerable interest, since they bear on our understanding of the early dynamics associated with crop species origins and divergence, the nature of human/plant interactions, and the genetic basis of domestication. Moreover, an understanding of the evolutionary history of crop species aids genetic mapping approaches, as well as informs plant breeding strategies.

Within the genus *Oryza*, crop domestication has occurred at least twice – once in Asia and separately in Africa. In Asia, the wild rice *O. rufipogon* was domesticated into the Asian rice *O. sativa* approximately 9,000 years ago, while independently in West Africa, the wild rice *O. barthii* was domesticated into the African rice *O. glaberrima* about 3,000 years ago [4]. Recent archaeological studies have also suggested that a third independent domestication event occurred in South America during pre-Columbian times, but this crop species is no longer cultivated [5].

The domestication history of Asian rice has been extensively studied both from the standpoint of archaeology [6] and genetics [7]. In contrast, much less is known about the domestication of *O. glaberrima*. Based on the morphology of rice grown in West Africa, the ethnobotanist Portères was the first to postulate an *O. glaberrima* domestication scenario [8,9], in which he hypothesized the inner Niger delta region in Mali as the center of domestication. He based this hypothesis on *O. glaberrima* in this area having wild rice-like traits (termed “genetically dominant characteristics” by Portères). In contrast, *O. glaberrima* with domesticated rice-like traits (termed “genetically recessive characteristics” by Portères) were found in two geographically separated regions: (i) the Senegambia region bordering the river Sine to the north and river Casamance to the south; and (ii) the mountainous region of Guinea. Portères hypothesized the derived traits observed in *O. glaberrima* from Senegambia and Guinea were due to those regions being secondary centers of diversification, but the inner Niger delta region remained as the primary center of diversity for African rice. Archaeological studies have found ceramic impressions of rice grains in north-east Nigeria dating ∼3,000 years ago, but documented *O. glaberrima* has only been found in the inland Niger delta at Jenne-Jeno, Mail dating ∼2,000 years ago [10].

Few population genetic studies have attempted to understand the evolutionary history and geographic structure of *O. glaberrima*. Microsatellite-based analysis showed genetic structure within *O. glaberrima* [11], suggesting the phenotypic differences observed by Portères may have stemmed from this population structure. With high-throughput sequencing technology, population genomic analysis indicated *O. barthii*, the wild progenitor of *O. glaberrima*, had evidence of population structure as well, dividing into 5 major genetic groups (designated as OB-I to OB-V). The OB-V group from West Africa was mostly closely related to *O. glaberrima*, suggesting that this *O. barthii* group from West Africa was likely to be the direct progenitor of African rice [12]. Genome-wide polymorphism data also indicated that *O. glaberrima* had a population bottleneck spanning a period of >10,000 years, indicating a protracted period of pre-domestication related management during its domestication [13].

While genome-wide variation studies have given valuable insights into the evolutionary history of *O. glaberrima*, they have not necessarily examined how *O. glaberrima* was domesticated from *O. barthii*. This is because the domestication history of a crop is best examined from the pattern of variation observed in genes underlying key domestication phenotypes [2]. Crop domestication accompanies a suite of traits, often called the domestication syndrome, which modified the wild progenitor into a domesticated plant that depends on humans for survival and dispersal [1]. In rice these traits include the loss of seed shattering [14,15], plant architecture change for erect growth [16,17], closed panicle [18], reduction of awn length [19,20], seed hull and pericarp color changes [21,22], change in seed dormancy [23], and change in flowering time [24]. During the domestication process, it is likely that these traits were not selected at the same time and selection would have occurred in stages depending on the importance of the domestication trait. Traits such as loss of seed shattering and plant erect growth would have been among the initial phenotypes humans have selected to distinguish domesticates from their wild progenitors, while traits that improved taste and appearance of the crop, or adaptation to the local environment would likely have been favored in later diversification/improvement stages of crop evolution [3,25].

Genes involved in the early stage domestication process are key to understanding the domestication process of a crop. Sequence variation from these early stage domestication genes can indicate whether a specific domestication trait had single or multiple causal mutations, revealing whether domestication had a single or multiple origins. Furthermore, the geographic origin and spread of domestication traits can be inferred from sequence variation in domestication loci within contemporary wild and domesticate populations [14,26–29]. In Asian rice, for example, genome-wide single nucleotide polymorphisms (SNPs) have suggested that each rice subpopulation had independent wild rice populations/species as their progenitors [30–34], but the domestication genes revealed a single common origin of these loci [32], suggesting a single *de novo domestication model for Asian rice [34–39]. On the other hand in barley*, the domestication gene for the non-brittle phenotype (*btr1* and *btr2*) had at least two independent origins [40,41], likely from multiple wild or proto-domesticated individuals

[42]. This suggests barley follows a multiple domestication model [43–45] originating from a genetically mosaic ancestral population [42,46,47].

To better understand the domestication of *O. glaberrima*, we have re-sequenced whole genomes of *O. glaberrima* landraces and its wild progenitor *O. barthii* from the hypothesized center of origin in the inner Niger delta, the middle and lower Niger basin that includes the countries Niger and Nigeria, and from Central Africa which includes Chad and Cameroon. The latter two regions were not heavily sampled in previous genomic studies. Together with published *O. barthii* samples from West Africa [12] and *O. glaberrima* samples from the Senegambia and Guinea region [13], we conducted a population genomic analysis to examine the domestication history of *O. glaberrima*, and an evolutionary analysis of genes involved in the early stage domestication process, mainly in the traits involving loss of shattering and erect plant growth,. With our data we examine the evolutionary and population relationships between *O. glaberrima* and *O. barthii*, the demographic history, and the geographic origin(s) of domestication of the African rice *O. glaberrima*.

## Results and Discussion

### Sequence diversity in *O. glaberrima* and *O. barthii*

We re-sequenced the genome of 80 *O. glaberrima* landraces from a geographic region that spanned the inner Niger delta and lower Niger basin region (S1A Fig). Together with 92 *O. glaberrima* genomes that were previously re-sequenced [13], which originated mostly from the coastal region (S1B Fig), the 172 *O. glaberrima* genomes analyzed in this study represent a wide geographical range from West and Central Africa. We also re-sequenced the genomes of 16 *O. barthii* samples randomly selected from this area (S1C Fig), and analyzed it together with 94 *O. barthii* genomes that were previously re-sequenced [12].

The average genome coverage in the data set we gathered for this study was ∼16.5× for both domesticated and wild African rice samples, and was comparable to the sequencing depth (∼16.1×) in our previous study. The Wang *et al*. [12] study sequenced a subset of their samples to a higher depth (∼19.4×), although the majority of their samples had relatively low coverage (∼3.9×) (see S1 Table for genome coverage of all samples in this study). To avoid potential biases in genotyping that arises from differences in genome coverage [48,49], we conducted our population genetic analysis using a complete probabilistic model to account for the uncertainty in genotypes for each individual [50,51]. A subset of our analysis, however, required hard-called genotypes, and SNPs were thus called from individuals with greater then 10× genome-wide coverage. After quality control filtering a total of 634,418 and 1,568,868 SNPs from *O. glaberrima* and *O. barthii* respectively were identified from non-repetitive regions.

### Genetic and geographic structure of *O. glaberrima*

The genetic structure across domesticated and wild African rice was examined by estimating the ancestry proportions for each individual in our dataset. We used the program NGSadmix [52], which uses genotype likelihoods from each individual for ancestry estimation and is based on the ADMIXTURE method [53]. Ancestry proportions were estimated by varying the assumed ancestral populations (K) as 2 to 9 groups (S2 Fig). With K=2 it divided the data set into *O. glaberrima* and *O. barthii* species groups, with several *O. glaberrima* samples having varying degrees of *O. barthii* ancestry (ranging from 4.5 to 40.5%). Interestingly, there were a number of *O. barthii* samples that had high proportions of *O. glaberrima* ancestry. All these wild rice with discernible *O. glaberrima* admixture corresponded to the OB-V *O. barthii* group designated in Wang *et al*. [12]; this group was hypothesized to be the progenitor of *O. glaberrima*. However, our ancestry analysis suggests that the close phylogenetic relationship of this wild *O. barthii* group could also be a result of wild-domesticated rice hybridization (see below) [54].

Increasing K further subdivided *O. glaberrima* into subpopulations. At K=5 there were three major genetic groups within *O. glaberrima* and two within *O. barthii* (Fig 1A left panel). The two *O. barthii* genetic groups corresponded to OB-I and OB-II group identified in Wang et al. [12]. For *O. glaberrima*, the ancestry proportions showed structuring into 3 major geographic regions: coastal, inner Niger delta, and lower Niger basin populations (Fig 1A right panel). At K=7 *O. glaberrima* divided into 5 genetic groups (Fig 1B left panel), where the coastal and inner Niger delta population identified at K=5 were divided into northern and southern genetic groups (Fig 1B right panel).

**Fig 1.**
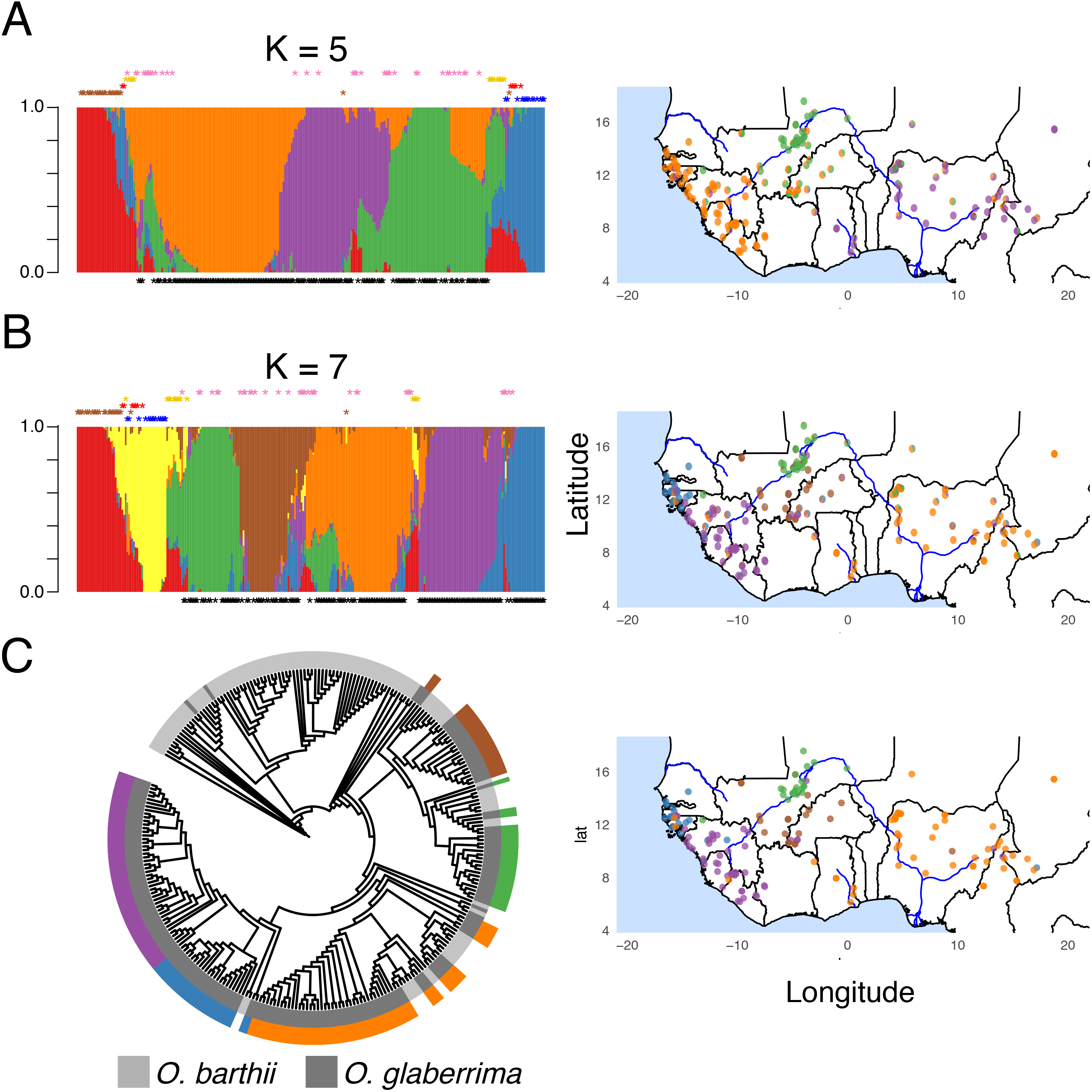
Population structure within *O. glaberrima*. (A) Ancestry proportions estimated from NGSadmix assuming K=5 and (B) K=7. Black stars below the admixture barplot indicate *O. glaberrima* individuals. Colored stars above admixture barplot are the *O. barthii* grouping designated by Wang et al. [12] where blue: OB-I, brown: OB-II, red: OB-III, yellow: OB-IV, and pink: OB-V group. (C) Neighbor-joining tree built using a distance matrix estimated from NGSdist. Color strips represent group of *O. glaberrima* individuals sharing a common ancestor. Geographical distribution of the ancestry proportion or phylogenetic grouping is seen in the right panel.

Phylogenomic analysis were then conducted using genotype likelihoods to estimate the pairwise genetic distances [55] and build a neighbor-joining tree (Fig 1C left panel). *O. glaberrima* formed a paraphyletic group with several *O. barthii* individuals. Consistent with the ancestry result at K=7, *O. glaberrima* was divided into 5 groups in the phylogeny. The geographic distribution of these 5 phylogenetic groups (Fig 1C right panel) were also concordant with the geographic distribution of the ancestry components in the African rice subpopulations identified at K=7.

The majority of the newly sequenced *O. barthii* from this study belonged to either OB-I or OB-II subpopulations designated by Wang *et al*. [12] (S3 Fig). The ancestry proportion for the OB-III and OB-IV groups suggested these individuals were an admixed group, with OB-III an admixture of OB-I and OB-II, and the OB-IV group possessing a mix of ancestry from both wild and domesticated rice. Unlike the OB-V group of *O. barthii*, however, which also had several individuals of mixed wild and domesticated rice ancestry, the OB-IV group did not phylogenetically cluster with *O. glaberrima* (Fig 1 and S2 Fig). This suggests that the OB-IV subpopulation may be an evolutionary distinct population, and the ancestry proportions were possibly mis-specified [56]. Hence, we considered individuals that were monophyletic with the OB-I or OB-II subpopulations as the wild *O. barthii* subpopulation and henceforth designated it as OB-W (Fig 2A). *O. barthii* that were paraphyletic with *O. glaberrima* were considered as a separate *O. barthii* group and designated as OB-G (Fig 2A). All other *O. barthii* not grouped as OB-W or OB-G were excluded from downstream analysis. Geographically, OB-G was found throughout West Africa but OB-W was found mostly in inland West African countries such as in Mali, Cameroon, and Chad (S2 Table).

**Fig 2.**
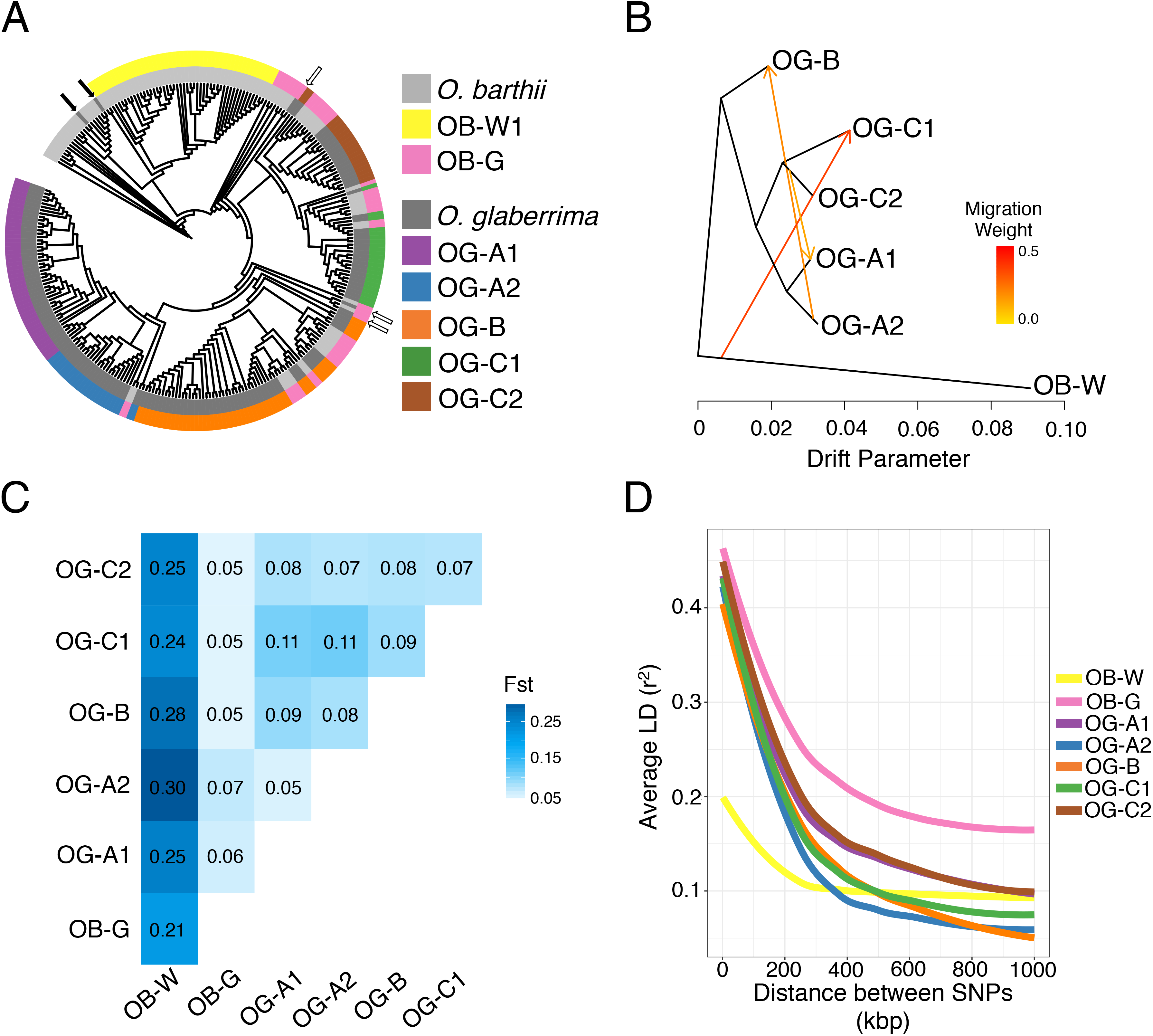
Evolutionary relationship between wild and domesticated African rice. (A) Neighbor-joining tree for all 282 samples analyzed in this study. Color strips represent group of individuals sharing a common ancestor and that were the focus of this study. Arrows indicate two *O. glaberrima* individuals clustering with *O. barthii*. (B) Treemix admixture graph of the 5 *O. glaberrima* subpopulations and the *O. barthii* outgroup. Arrows represent admixture between populations. (C) Pairwise Fst values between *O. glaberrima* and *O. barthii* genetic groups. (D) Average linkage disequilibrium between pair of SNPs.

In *O. glaberrima*, it appears there are at least 5 population groups (Fig 2A). Consistent with the hypothesized Guinea highland and Senegambia populations, the coastal populations were divided into OG-A1 and OG-A2 genetic groups (collectively the OG-A supergroup). The lower Niger basin and central African individuals formed as a single OG-B group. Finally, for the inner Niger Delta region, landraces closest to the delta formed the OG-C1 group while the others formed the OG-C2 group; collectively these represent the OG-C supergroup. There were 3 *O. glaberrima* individuals (IRGC104883, IRGC105038, and IRGC75618) that were not forming a monophyletic cluster with the 5 population groups (Fig 2A white arrow), but rather formed as a sister group to all *O. glaberrima* or sister group to both OG-A and OG-B group. Because they were most closely related to OB-G samples we considered them as OB-G as well.

Interestingly, there were two *O. glaberrima* individuals (IRGC103631 and IRGC103638) that phylogenetically clustered with *O. barthii* (Fig 2A filled arrows). Ancestry estimates for the two samples showed high proportions of both *O. glaberrima* and *O. barthii* ancestry (S4 Fig). Because their ancestry profiles were similar to the OB-IV group, this suggested IRGC103631, IRGC103638, and the OB-IV group rice might represent feralized rice with mixed wild and domesticated ancestry. These two *O. glaberrima* samples were not used in subsequent analysis.

### Evidence of admixture within *O. glaberrima*

The ancestry proportions (Fig 1A and B) suggested several individuals were admixed and these would not be properly represented in the neighbor-joining tree (Fig 2A). To estimate gene flow, we inferred relationships between genetic groups as admixture graphs using Treemix [57], rooting the graph with the OB-W group. Without modeling migration events, Treemix analysis suggests a different topology compared to the neighbor-joining tree, where the OG-A and OG-C were sister groups while OG-B as the outgroup to OG-A/OG-C (S5A Fig). This model explains 96.9% of the variance. Treemix models sequentially fitting an increasing number of migration edges did not change the topology, but improved the model fit (S5B and S5C Figs) and with 3 migration edges almost all of the variance could be explained (Fig 2B). There were two migration events between *O. glaberrima* genetic groups: (i) OG-B traced 20.1% of its ancestry to OG-A2, and (ii) OG-A1 traced 13.5% of its ancestry to OG-C2. There was also evidence of admixture between *O. barthii* and *O. glaberrima* where 36.1% of the ancestry for OG-C1 was derived from an ancient OB-W-like population. Consistent with the migration edge detected between OB-W and OG-C1 by Treemix, a four-population test [58] of the tree [[OB-W,OG-B],[OG-C1,OG-C2]] showed significant deviation from tree-ness (Z-score=5.26), suggesting admixture between OB-W and OG-C1; or OG-B and OG-C2 genetic groups.

As an alternative method of examining admixture, we calculated the D-statistics across the four population H1, H2, H3, and H4, testing for evidence of gene flow between H3 and either H1 or H2 [59]. Extended D-statistics were implemented to analyze both low and high coverage data for multiple individuals together [60]. D-statistics largely corroborated the Treemix results, with significant admixture between OG-C2 and both OG-A genetic groups; and significant admixture between OG-A2 and OG-B (Table 1). None of the D-statistics testing admixture between *O. glaberrima* and OB-W were significant (S3 Table). It is possible the D-statistic did not detect the OB-W and OG-C1 gene flow, if the migration involved a wild *O. barthii population related to OB-W. This* may partially explain the wild rice-like traits Portères had observed from *O. glaberrima* in the inner Niger delta [8].

**Table 1.** D-statistics for four populations (H1, H2, H3, H4).

| **D** | **Z** | **nABBA** | **nBABA** | **H1** | **H2** | **H3** | **H4** |
| --- | --- | --- | --- | --- | --- | --- | --- |
| -0.090 | **-4.92** | 75614.98 | 90512.23 | OG-C2 | OG-C1 | OG-A1 | *O. rufipogon* |
| -0.090 | **-5.82** | 75619.17 | 90519.52 | OG-C2 | OG-C1 | OG-A2 | *O. rufipogon* |
| 0.029 | 1.69 | 71535.48 | 67522.61 | OG-A1 | OG-A2 | OG-B | *O. rufipogon* |
| -0.039 | -2.59 | 81210.78 | 87823.07 | OG-C2 | OG-C1 | OG-B | *O. rufipogon* |
| -0.006 | -0.32 | 68973.69 | 69769.31 | OG-A1 | OG-A2 | OG-C1 | *O. rufipogon* |
| -0.006 | -0.28 | 69398.97 | 70184.05 | OG-A1 | OG-A2 | OG-C2 | *O. rufipogon* |
| Topology assuming OG-A and OG-B group are sister group |  |  |  |  |  |  |  |
| -0.042 | -2.08 | 80866.84 | 87896.26 | OG-A1 | OG-B | OG-C1 | *O. rufipogon* |
| -0.037 | -1.98 | 79552.37 | 85743.22 | OG-A2 | OG-B | OG-C1 | *O. rufipogon* |
| -0.091 | **-4.60** | 76537.12 | 91854.87 | OG-A1 | OG-B | OG-C2 | *O. rufipogon* |
| -0.088 | **-5.04** | 75290.69 | 89812.19 | OG-A2 | OG-B | OG-C2 | *O. rufipogon* |
| Topology assuming OG-A and OG-C group are sister group |  |  |  |  |  |  |  |
| 0.029 | 1.69 | 71535.48 | 67522.61 | OG-A1 | OG-A2 | OG-B | *O. rufipogon* |
| -0.028 | -1.32 | 76537.12 | 80941.84 | OG-A1 | OG-C2 | OG-B | *O. rufipogon* |
| -0.064 | -3.05 | 80866.84 | 91885.78 | OG-A1 | OG-C1 | OG-B | *O. rufipogon* |
| -0.053 | -2.47 | 75290.69 | 83731.72 | OG-A2 | OG-C2 | OG-B | *O. rufipogon* |
| -0.086 | **-4.06** | 79552.37 | 94585.89 | OG-A2 | OG-C1 | OG-B | *O. rufipogon* |
| -0.039 | -2.59 | 81210.78 | 87823.07 | OG-C2 | OG-C1 | OG-B | *O. rufipogon* |

D, ABBA-BABA D-statistics; Z, Z-scores higher then |3.9| (p-value <9.6 × 10^‒5^) are bolded; nABBA and nBABA, Sites that had ABBA or BABA configuration respectively; H1,H2,H3,H4, Four populations with phylogenetic relationships where H1 and H2 population are sister groups, H3 is an outgroup to both H1 and H2 population, and H4 is the outgroup to H1, H2, and H3 population.

### The OB-G group of *O. barthii* is possibly a feral African rice

Previous molecular studies have argued the close genetic affinities of some west African *O. barthii* (namely the OB-G group in this study) to *O. glaberrima*, as evidence of the former being the progenitor population of African rice [12,61]. We thus examined the properties of the OB-G group in relation to OB-W and *O. glaberrima*.

First, we found that the level of population differentiation between OB-G and *O. glaberrima* was low (Fig 2C), almost comparable to the level seen between *O. glaberrima* genetic groups. In contrast, there is a modest level of differentiation between each *O. glaberrima* genetic group and OB-W (Fst ∼ 0.26), and the OB-G group also had similar high levels of differentiation to OB-W (Fst ∼ 0.21).

Second, we examined levels of linkage disequilibrium (LD) decay, as wild and domesticated populations have different LD profiles, due to the latter undergoing domestication-related bottlenecks and selective sweeps [62]. In the African rice group, as expected, all *O. glaberrima* genetic groups had higher levels of LD compared to the OB-W group (Fig 2D). The OB-G group also had high levels of LD that was comparable to those observed in *O. glaberrima*. Genome-wide polymorphism levels for the OB-G group were also comparable between OB-G and *O. glaberrima*. Specifically, compared to the OB-W group, SNP levels and Tajima’s D [63] were significantly lower in both OB-G and *O. glaberrima* (S6 Fig).

Third, we examined the demographic histories of the *O. barthii* groups and *O. glaberrima*. PSMC’ [64] was then used to estimate past demographic changes for each genetic group; domestication often leads to bottlenecks [13,65,66]. Because African rice is expected to be a predominant selfer, we created pseudo-diploid genomes [67] for all pairwise combinations within a genetic group. The results showed two different patterns. OB-W is composed of two genetic groups (Fig 2A), and pseudo-diploids comparisons between members of the two groups, including those from Meyer *et al*. [13], did not exhibit a protracted bottleneck (S7 Fig). Instead, the PSMC’ plot showed a rapid increase in population sizes ∼ 30 kya (Fig 3A), likely due to the lineage splitting event between the two individuals that was used to generate the pseudo-diploid [68]. In contrast, several comparisons among individuals in the OB-W group has PSMC’ profiles that resembled those seen in *O. glaberrima*, suggesting a subset of the OB-W group may have undergone a population bottleneck as well. Moreover, PSMC’ profiles showed the demographic history of OB-G was largely concordant with those observed in *O. glaberrima* (Fig 3A). This bottlenecking in *O. barthii* suggests changes in the environment may have initiated the protracted bottleneck across *Oryza* in West Africa, possibly caused by the gradual desertification in the Sahara region [69]. It also further highlights similarities in the demographic histories of the OB-G group and *O. glaberrima*.

**Fig 3.**
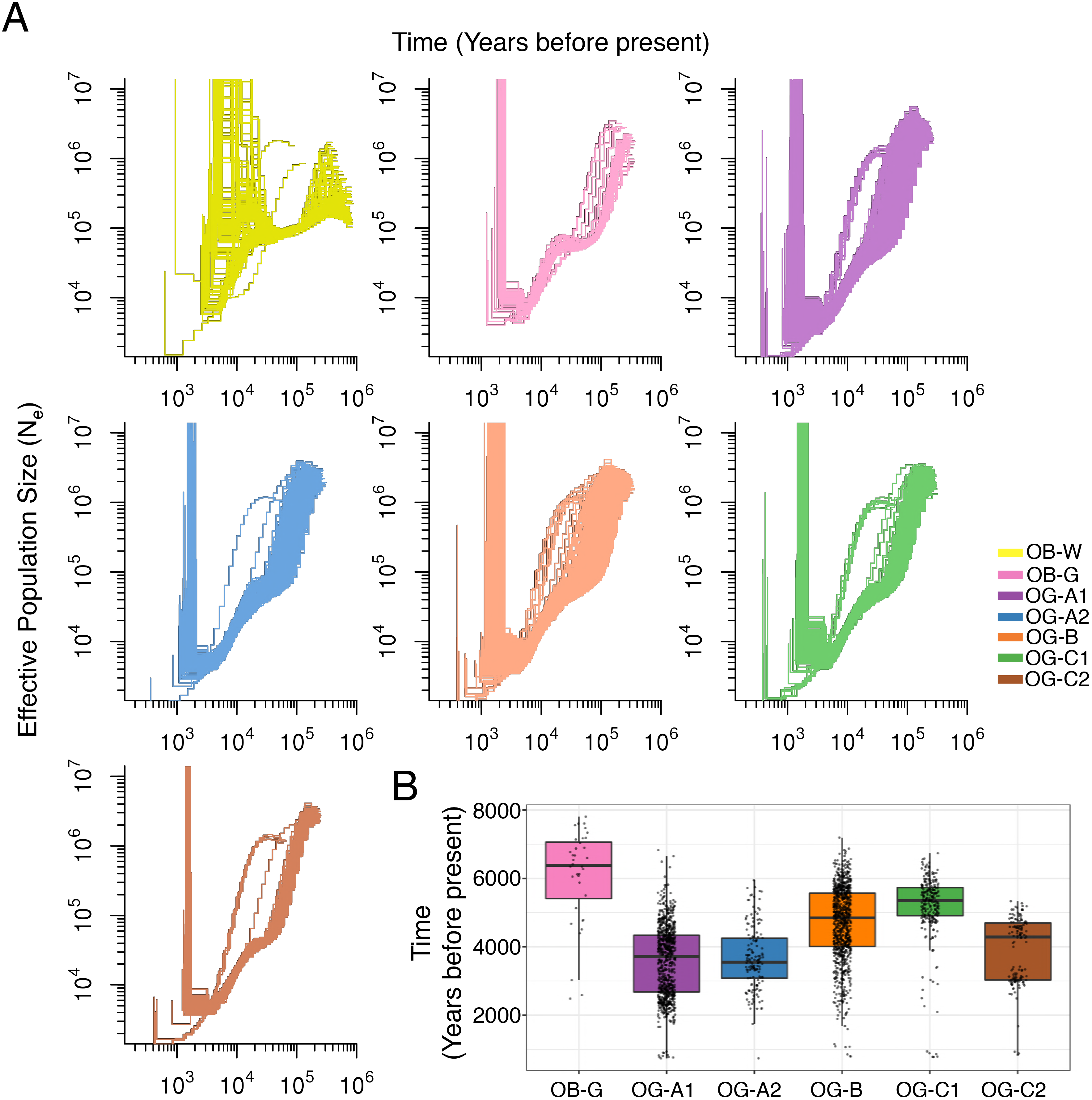
Demography of *O. glaberrima* and *O. barthii*. (A) PSMC’ estimated demography changes in *O. glaberrima* and *O. barthii* genetic groups. (B) Time at lowest population size in *O. glaberrima* and OB-G genetic group.

Together, the levels of genetic differentiation, linkage disequilibrium, SNP levels and patterns, and demographic profiles all suggest that OB-G behaves more like *O. glaberrima* than *O. barthii*. Furthermore, the majority of the OB-G samples carried at least one domestication mutation (see domestication gene haplotype analysis section for detail), further calling into question its status as the wild progenitor. In contrast all OB-W individuals do not carry the causal mutation/deletion at known domestication genes. Together, this suggests the genetic affinities were caused because many of them may represent feral weedy rice [70], resulting from the hybridization of domesticated and wild African rice. These intermediate type rice were found in a wide geographic region along the Niger river [71], and our study shows that most of them carry causal domestication mutations. This shows that OB-G may have formed after the domestication event and supports a de-domestication (endoferality) origin for that group [54]. Because of these observations, we caution future *O. glaberrima* domestication studies as these feral rice can lead to spurious domestication models.

### Elucidating the domestication origin of *O. glaberrima:* Contrasting demographic histories

A closer examination of the PSMC’ profiles in each *O. glaberrima* genetic group showed evidence of a protracted period of population size reduction, which had been previously observed in the coastal population [13]. However, the time when the population size reached its lowest level was different for each genetic group (Fig 3B). Specifically, the inland genetic groups reached their lowest effective population sizes (N_e_) on average 4.3 to 5.3 kya, while the costal genetic groups reached their lowest effective population sizes on average 3.5 to 3.7 kya, suggesting domestication bottleneck may be older in the inland groups than those found in the coast. Although this is consistent with the Portères domestication model, it is equally possible coastal populations had a longer domestication or cultivation period.

### The geographic origin of the *sh4* nonsense allele

To further identify the domestication origin(s) of *O. glaberrima*, we examined the haplotypes for the domestication genes involved in the non-shattering phenotype (*sh4* and *sh1*) and erect plant growth (*PROG1*) in both wild and domesticated African rice. We took an approach we term *functional phylogeography*, where we examined the haplotype structure surrounding the domestication gene of interest, inferred a haplotype phylogenetic network, and determined the geographic origin and spread of the functional mutation by comparing the geographic distributions of haplotypes in wild and domesticated African rice in a phylogenetic context.

For the *sh4* gene (*O. glaberrima* chromosome 4:25,150,788–25,152,622) the causal domestication mutation has been identified, and the gene also had evidence of a selective sweep [72,73]. The haplotype structure around the *sh4* gene showed most of the *O. glaberrima* landraces carried the causal domestication mutation, a C-to-T nonsense mutation at chromosome 4 position 25,152,034 that leads to a loss-of-function allele (Fig 4A arrow). Several individuals within OG-A1 group, including the reference genome, did not carry the causal mutation but still had long tracks of homozygosity at the *sh4* locus (Fig 4A star). A four-gamete test [74] of a region 4 kbp upstream, 2 kbp downstream, and spanning the *sh4* gene detected evidence of recombination, but it was only found within the *O. barthii* population (both OB-G and OB-W) and not within *O. glaberrima*. A neighbor-joining tree of the same region showed all *O. glaberrima* populations were divided into two major phylogenetic groups (S8 Fig). *O. glaberrima* individuals without the causal mutation (Fig 4A star) formed their own phylogenetic group (S8 Fig star).

**Fig 4.**
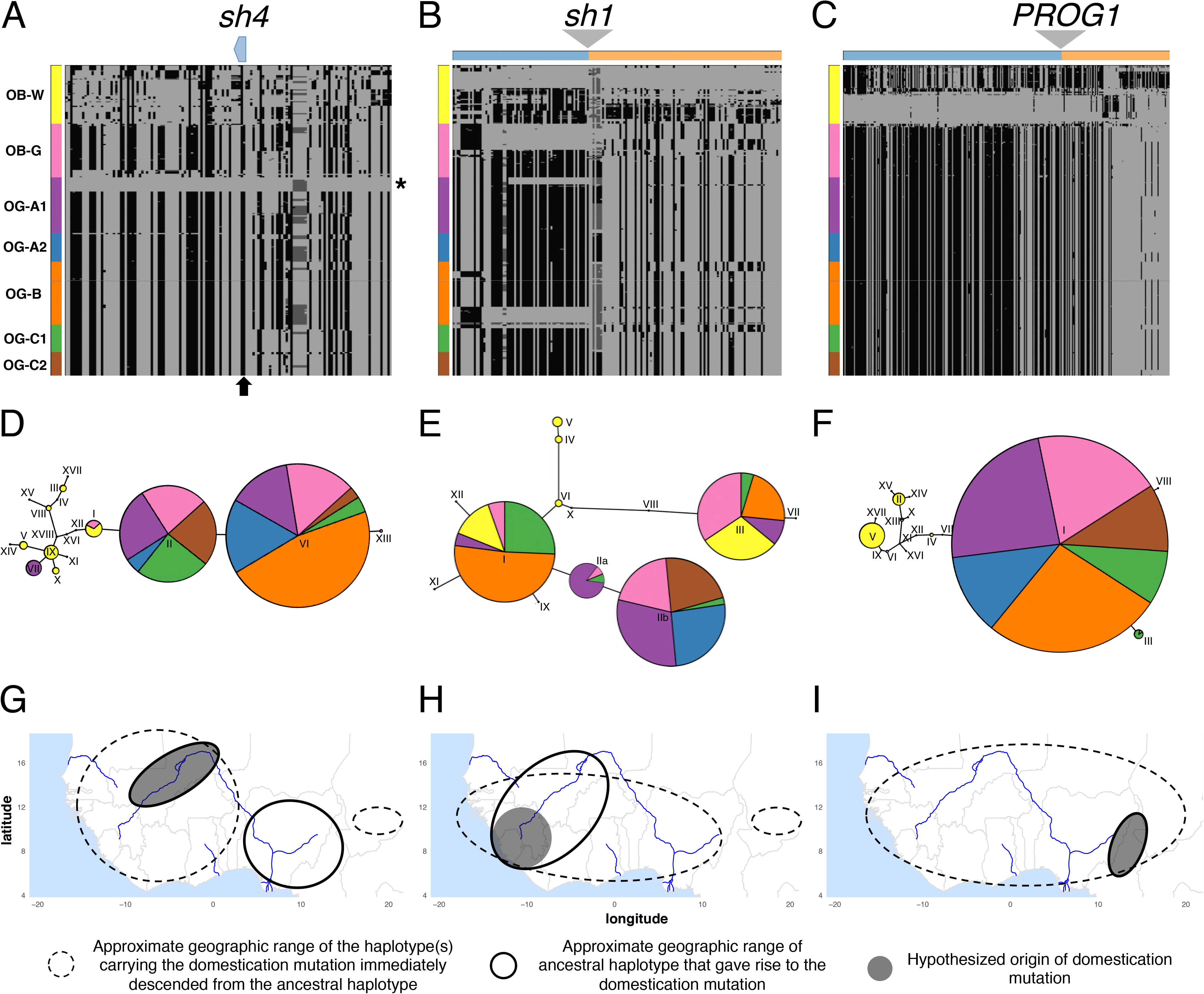
Haplotype analysis of the three domestication genes (A,D,G) *sh4*, (B,E,H) *sh1*, and (C,F,I) *PROG1*. Haplotype structures are shown for the genes (A) *sh4*, (B) *sh1*, and (C) *PROG1*. Homozygote genotype not identical to reference genome is shown in dark grey, heterozygote genotype shown in lighter shade of grey, and homozygote genotype identical to reference genome shown in lightest shade of grey. Regions are showing polymorphic sites from 25 kbp up- and downstream of the domestication gene. In *sh4* (A) the position of the causal domestication mutation is shown in arrow and OG-A1 samples without the causal mutation are indicated with a star. In *sh1* (B) and *PROG1* (C) region upstream and downstream the deletion are color coded above the haplotype structure. Haplotype network are shown for genes (D) *sh4*, (E) *sh1*, and (F) *PROG1*. Approximate geographic origins for the causal domestication mutation haplotype and its most closely related ancestral haplotype are shown for genes (G) *sh4*, (H) *sh1*, and (I) *PROG1*. See text for discussion of the hypothesized geographic origins.

To determine the origin of the non-shattering trait, we reconstructed the haplotype network of the non-recombining region of the *sh4* gene in all *O. glaberrima* and *O. barthii* genetic groups (Fig 4D). Majority of the *O. glaberrima* and OB-G group *sh4* haplotypes belonged to haplotypes II, VI, and XIII and they all shared the nonsense mutation. The two main haplotypes II and VI corresponds to the difference observed in the upstream region of the *sh4* gene (Fig 4A), with haplotype II evolving prior to haplotype VI. The closest haplotype to II was haplotype I, which was separated by two mutations (position 25,146,871 and the causal domestication mutation 25,152,034).

We tabulated the geographic distributions of *O. glaberrima* haplotypes II and VI/XIII, and haplotype I from the *O. barthii* OB-W group (Table 2). The ancestral haplotype I is found in 13 *O. barthii* individuals (4 OB-G group and 9 OB-W group), and these individuals originated over a wide geographic region of West Africa that includes both coastal and inland areas (See S4 Table for full list of members of each haplogroup and their country of origin). Of those in OB-W, 2 are from Mali, 2 from Nigeria and 5 are from Cameroon. Among the *O. glaberrima* that have the *sh4* mutation, the older haplotype II is found mostly in Mali, Burkina Faso and also Guinea.

**Table 2.** Geographical of origin for domestication gene haplotypes identified in Fig 4.

| Country of Origin | *sh4* |  |  |  | *sh1* |  |  |  |  | *PROG1* |  |  |  |  |  |
| --- | --- | --- | --- | --- | --- | --- | --- | --- | --- | --- | --- | --- | --- | --- | --- |
|  | others | I (OB-W) | II * | VI/XIII * | others | I | IIa | IIb * | III | others | XI | XII | IV | VII * | I/VIII/III * |
| Senegal | - | - | 3 | 12 | 3 | 1 | 1 | 15 | 3 | - | - | - | - | - | 19 |
| Gambia | - | - | - | 1 | - | 1 | - | 3 | - | - | - | - | - | - | 3 |
| Guinea-Bissau | - | - | 1 | 7 | - | 1 | 1 | 6 | - | - | - | - | - | - | 8 |
| Guinea | 2 | - | 10 | 8 | 4 | 3 | 11 | 8 | 2 | 1 | - | 1 | - | - | 23 |
| Sierra Leone | 1 | - | 5 | 4 | 2 | 2 | 6 | 2 | 1 | 1 | - | - | - | - | 11 |
| Liberia | 5 | - | 1 | 2 | - | - | 3 | 5 | - | - | - | - | - | - | 8 |
| Cote d'Ivoire | 2 | - | 1 | 2 | 2 | 1 | - | 2 | 2 | - | - | - | - | - | 5 |
| Ghana | - | - | - | 5 | - | 5 | - | - | - | - | - | - | - | - | 5 |
| Burkina Faso | - | - | 10 | 2 | 1 | - | 2 | 11 | 1 | - | - | - | - | - | 14 |
| Mali | 1 | 2 | 27 | 5 | 18 | 25 | 1 | 14 | 9 | 15 | - | - | - | - | 25 |
| Niger | - | - | - | - | 1 | 1 | - | - | 1 | 1 | - | - | - | - | 1 |
| Nigeria | - | 2 | - | 32 | 21 | 21 | - | 1 | 20 | 3 | - | - | - | - | 40 |
| Cameroon | - | 5 | - | 8 | 16 | 6 | - | - | 14 | 8 | - | - | 2 | 1 | 11 |
| Chad | - | - | 1 | 7 | 15 | 7 | - | 2 | 10 | 13 | 1 | - | - | - | 10 |
| Tanzania | - | - | - | - | 1 | - | - | - | - | 1 | - | - | 1 | - | - |
| Zambia | - | - | - | - | 1 | - | - | - | - | 1 | - | - | - | - | - |
| Botswana | - | - | - | - | 1 | - | - | - | - | 1 | - | - | - | - | - |
| Unknown | - | - | - | - | 1 | - | - | 4 | 1 | - | - | - | - | - | 5 |

\*Haplotypes carrying the domestication mutation. For haplotypes without the casual domestication mutation, only those that are most closely related to the haplotype with the causal mutation are shown. Haplotype numbers and their corresponding relationships are shown in Fig 4.

Here, we make the assumption that the areas of overlap between the ancestral haplotype (without the causal mutation) and the derived haplotype (with the causal mutation) is likely the place of origin of the domestication allele. For *sh4*, the distribution of haplotype II overlaps with haplotype I in Mali, pointing to Mali as being a likely place of origin for the *sh4* nonsense mutation (Fig 4G). The haplotypes VI and XIII thus subsequently evolved from haplotype II, which expanded over a much wider area, particularly in the Senegambia, and also to Nigeria, Cameroon and Chad. It should be noted that the sample size for haplotype I among OB-W is relatively small (n=9) leading to disjoint geographic ranges for its distribution (Fig 4G); therefore localizing the origin of the *sh4* causal mutation to Mali may be revised as more *O. barthii* samples are analyzed. However, haplotype II is found at highest frequency in Mali as well (∼46%, see Table 2), which provides further support for a Malian origin of the *sh4* mutations.

Wu *et al*. [73] had first noticed that several *O. glaberrima* individuals in the coastal region of West Africa did not have the causal domestication mutation in the *sh4* gene (Fig 4A star). Our data shows that all inland *O. glaberrima* carries the haplotype with the nonsense mutation, and the haplotype without the nonsense mutation was indeed limited to the coastal region, specifically in the OG-A1 genetic group. The haplotype network and neighbor-joining tree suggests these individuals had distinct evolutionary histories for the sh4 gene (Fig 4D and S8 Fig); they carry haplotype VII which is confined to Guinea. It remains to be determined whether the haplotype without the nonsense mutation can also result in a non-shattering phenotype. But assuming OG-A1 individuals are all non-shattering, the polymorphic status of the nonsense mutation suggests there has been an independent domestication for non-shattering in several OG-A1 individuals.

### The *sh1* gene deletion is polymorphic and has coastal origins

Assembly of the reference *O. glaberrima* genome had first shown the gene *sh1* was missing in the *O. glaberrima* genome but not in the *O. barthii* genome [12]. Recently, the *sh1* gene (see Materials and Method section “Shattering gene nomenclature” for comment on gene nomenclature of *sh1*) was identified as another causal gene for the non-shattering trait, and the causal mutation was indeed a gene deletion that was polymorphic in several coastal O. glaberrima populations (Lu *et al*. 2018. in press). A read depth based analysis (see Materials and Method section “Gene deletion analysis of genes *sh1* and *PROG1*” for details) showed the *sh1* gene was missing in several *O. glaberrima* individuals (Table 3). Specifically, no individuals in the inland genetic groups OG-B and OG-C1 had the *sh1* deletion, but in another inland genetic group OG-C2 all individuals carried the *sh1* deletion. In the coastal population all individuals from the Senegambia genetic group OG-A2 had the *sh1* deletion, while in the genetic group OG-A1 the deletion was polymorphic (frequency ∼ 32%). The deletion was also polymorphic in the OB-G group (frequency ∼ 33%) but no individual had a deletion in the OB-W wild group.

**Table 3.**
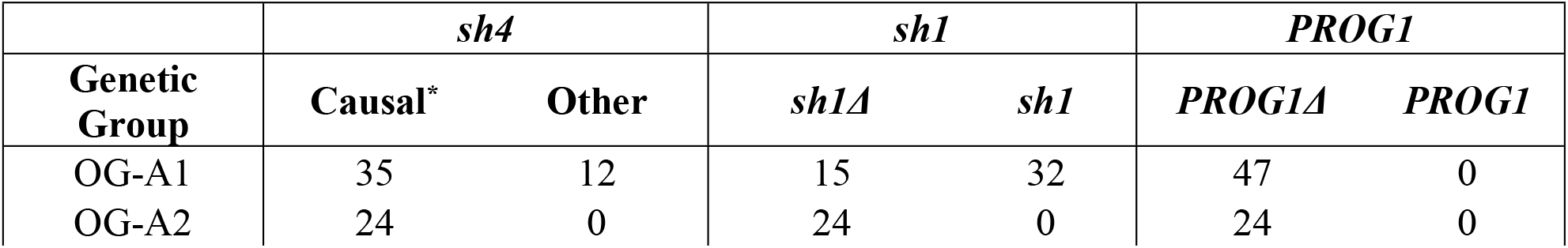
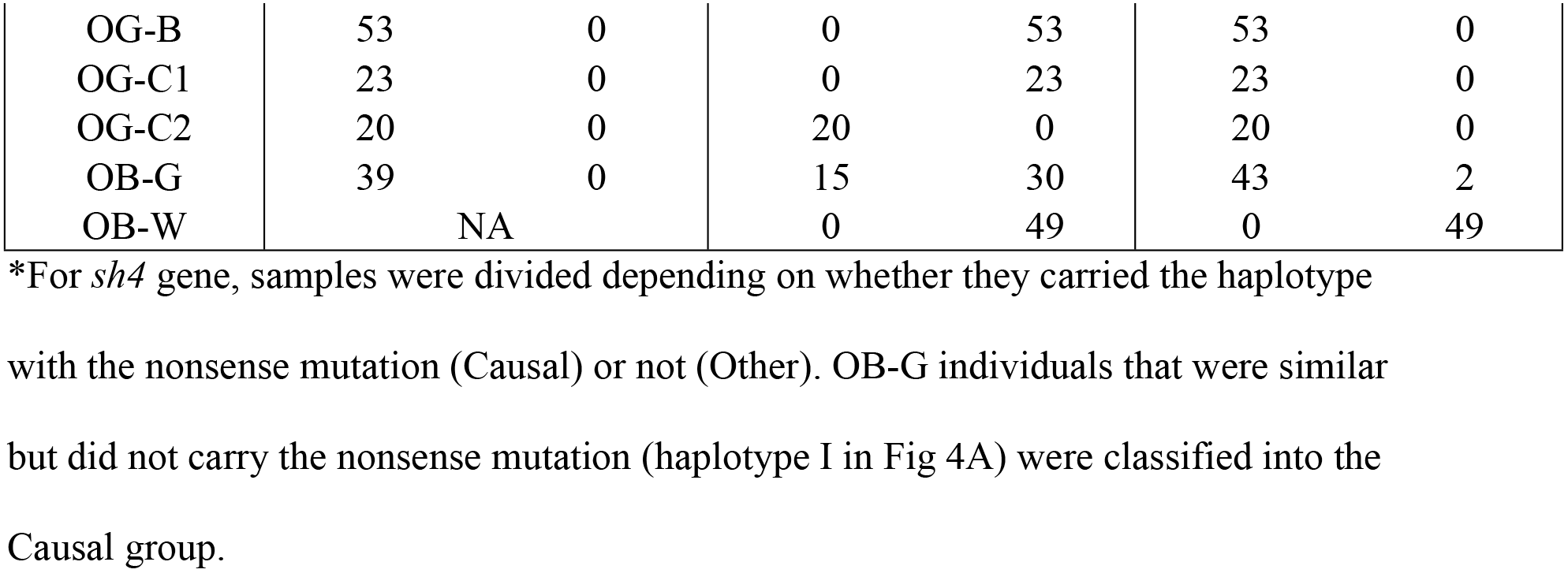
Domestication allele status in *O. glaberrima* and *O. barthii* groups.

Reflecting the polymorphic status of the *sh1* deletion, we observed no single haplotype at high frequency (Fig 4B). There was evidence of recombination in the downstream 5 kbp of the deletion in both wild and domesticated African rice, so an unambiguous haplotype network could not be inferred for that region. The haplotype network of a non-recombining 5 kbp region immediately upstream of the deletion with 22 polymorphic sites, indicated that there were 3 main *O. glaberrima* haplotypes – I, II and III (Fig 4E). All *O. glaberrima* in haplotypes I and III do not carry the *sh1* deletion (See S5 Table for full list of members of each haplogroup and their country of origin). Haplotype II can be further divided into two depending on the status of the *sh1* gene deletion. Haplotype IIb contains all of the *O. glaberrima* individuals with the *sh1* deletion, while haplotype IIa is found 23 OG-A1, 1 OB-G, and 1 OG-C1 individuals and does not carry the *sh1* deletion. A neighbor-joining tree of the upstream 5 kbp region confirmed the haplotype network, where all OG-A2 and OG-C2 individuals that had the deletion grouped together with several OG-A1 individuals (S9 Fig).

Together, these results indicate that the *sh1* deletion might have arisen on a genetic background most closely related to the OG-A1 genetic group, which in turn suggests a coastal origin for the *sh1* deletion. This coastal origin is supported by the geographic distributions of the different *sh1* haplotypes. Haplotype IIb, which contains the *sh1* deletion, is found across a wide area in West and Central Africa (Table 2). Haplotype IIa, which does not have the deletion and is presumably ancestral to IIb, is found at highest frequency in Guinea and Sierra Leone. Given where the distributions of haplotypes IIa and b overlap, it would suggest that a region encompassing the coastal countries of Guinea and Sierra Leone was where the *sh1* deletion originated (Fig 4H).

Another study (Lu et al. 2018. in press) had found the *sh4* and *sh1* double mutant was most prevalent in the Senegambia *O. glaberrima* individuals (identified here as the OG-A2 genetic group). Here, we corroborate their findings and further identify that the double mutant was also selected in the inland region but only in the OG-C2 genetic group (Table 3), which is found in southern Mali and Burkina Faso region (Fig 1). It is unclear why the double mutant had not spread further inland or existed in a polymorphic state in the OG-A1 genetic group, but aspects of this behavior is also seen in *O. sativa* where the causal mutation in the *qSh1* gene, which causes the non-shattering phenotype, is found only in the temperate japonica subpopulation [15]. Non-shattering is often considered as the trait selected in the earliest stages of rice domestication; however the process may have continued well into the diversification/improvement phase as well. In *O. glaberrima, sh1* and *sh4* double mutant causes a complete ablation of the abscission layer leading to a complete non-shattering phenotype (Lu *et al*. 2018. in press), which would have led to a higher yield per plant but at the cost of increased labor for separating the grains off the rachis [75]. Hence, the cost involved in non-shattering may have led to different preferences for the trade-off between harvesting and threshing among the people cultivating *O. glaberrima*, which could account for the polymorphism in the frequency of the mutation conferring non-shattering in both *sh4* and *sh1* genes.

### *PROG1* is deleted in all *O. glaberrima* and likely originated from central Africa

Analysis of *O. sativa* gene orthologs in *O. glaberrima* and *O. barthii* indicates that the gene *PROG1* is missing only in *O. glaberrima* (S6 Table). Synteny of the genes immediately surrounding *O. sativa PROG1* was maintained in both *O. glaberrima* and *O. barthii*, suggesting the *PROG1* gene is deleted specifically in *O. glaberrima*. Mutation in the *PROG1* gene causes *O. sativa* to grow erect which is hypothesized to increase growing density and enhance photosynthesis efficiency for increased grain yield [16,17]. Hence we examined the population genetics of the *PROG1* gene in *O. glaberrima* and *O. barthii* as a candidate domestication gene.

We conducted genome-wide sliding window analysis of the ratio of polymorphism in the wild OB-W group to all domesticated *O. glaberrima* (π_w_/ π_D_). A domestication-mediated selective sweep would lead to a reduction in nucleotide variation around the target domestication gene, but only within the domesticated group. Because *PROG1* is deleted in *O. glaberrima,* the selection signal will be around the candidate deletion region. Within 10 kbp of the candidate deletion region π_w_/ π_D_ was within the top 1% value, and this was observed regardless of whether the *O. glaberrima* or *O. barthii* genome was used as the reference genome in SNP calling (S10 Fig). This was consistent with the *PROG1* region having gone through a selective sweep during *O. glaberrima* domestication.

The sequencing read depth analysis found *PROG1* was missing in all *O. glaberrima* landraces (Table 3). None of the OB-W group had the *PROG1* deletion and all but two of the OB-G individuals had the *PROG1* deletion. We note this is the first candidate domestication gene that has been identified where the causal mutation is fixed in all *O. glaberrima* population. Polymorphisms surrounding the deletion comprised a single unique haplotype segregating across all *O. glaberrima* samples and most of the OB-G samples (Fig 4C). A haplotype network of a non-recombining 5 kbp region immediately upstream of the deletion showed that all individuals with the deletion belonged to the same major haplotype group, with the dominant haplotype I, as well as peripheral haplotypes VII, VIII, and III (Fig 4F and see S7 Table for full list of members of each haplogroup and their country of origin). Neighbor-joining tree of this region collapsed all *O. glaberrima* into a single phylogenetic group (S11 Fig), which suggests a single origin for the deletion.

Haplotype VII is the earliest haplotype with the deletion, and is found in an OB-G individual found in Cameroon. The ancestral haplotype IV is most closely related to all haplotypes with the deletion, and they consisted of three OB-W individuals: IRGC103912 (Tanzania), IRGC105988 (Cameroon), and WAB0028882 (Cameroon). The downstream region of the deletion was consistent with what is observed in the upstream region (S12 Fig and S8 Table for full list of members of each haplogroup and their country of origin), where 22 polymorphic sites from a non-recombining 7 kbp downstream region indicated the same OB-W individuals IRGC105988 and WAB0028882, both from Cameroon, were the most closely related haplotype to the *PROG1* deletion haplotype. Neighbor-joining trees of both the upstream and downstream regions also showed these two individuals to be the sister group to all *O. glaberrima* samples.

Together, the geographic distribution of the *PROG1* region haplotypes suggest that the *PROG1* deletion may have occurred in a wild progenitor closely related to those found in Cameroon in central Africa, and spread throughout West Africa through admixture with local *O. glaberrima* genetic groups (Fig 4I). Like in the case of the *sh4* phylogeographic analysis, the *PROG1* conclusion must be tempered by an acknowledgement that the sample size of ancestral haplotypes is small (n = 3).

Interestingly, a similar observation has been made in *O. sativa* where all Asian rice subpopulations are monophyletic in the *PROG1* region, but genome-wide the different Asian rice variety groups/subspecies do not share immediate common ancestors [32,76]. Recent studies show the importance of admixture during the domestication processes of crops [34,77–80], and in *O. glaberrima* it may have played a critical role in the fixation of this core domestication gene.

## Conclusion

Our analysis of whole genome re-sequencing data in the African rice *O. glaberrima* and its wild ancestor *O. barthii* provides key insights into the geographic structure and nature of domestication in crop species. Our analysis suggests that *O. glaberrima* is comprised of 5 distinct genetic groups, which are found in different geographic areas in West and Central Africa. There has been extensive gene flow between several of these genetic groups, which may have facilitated the spread of key domestication genes. Moreover, we find that many individuals that have been identified as *O. barthii* (and which in the past have been thought to be the immediate ancestor of the domesticated crop) form a distinct genetic group that behaves almost identically to *O. glaberrima*, including similarities in LD decay and demographic histories, and low genetic differentiation with domesticated African rice. Moreover, several of these *O. barthii* individuals carried causal mutations in the key domestication genes *sh4, sh1* and *PROG1*. Together suggests that these *O. barthii* individuals, which collectively we refer to as the OB-G group, may represent a feral *O. glaberrima* or may have been misidentified as the crop species.

Portères hypothesized that western inland Africa near the inner Niger delta of Mali as the center of origin for *O. glaberrima* [8,9], and this has been the commonly accepted domestication model for *O. glaberrima* [81]. Under this single center of origin model, *O. glaberrima* from the OG-C1 genetic group (closest to the inner Niger delta) would have acquired key domestication mutations before spreading throughout West Africa. Here, we suggest that the domestication of *O. glaberrima* may be more complex. Phylogeographic analysis of three domestication loci indicates that the causal mutations associated with the origin of *O. glaberrima* may have arisen in three different areas.

An alternative view suggests domestication has largely been a long protracted process, often involving thousands of years of transitioning a wild plants into a domesticated state [75,82,83]. If this indeed happened for *O. glaberrima*, our study suggests this protracted period of domestication had no clear single center of domestication in African rice. Instead domestication of African rice was likely a diffuse process involving multiple centers [83–85]. In this model, cultivation may have started at a location and proto-domesticates spread across the region with some (but maybe not all) domestication alleles. Across the multiple regions, the differing environmental conditions and cultural preferences of the people domesticating this proto-*glaberrima* resulted in differentiation into distinct genetic groups. Temporal and spatial variation in the domestication genes resulted in causal mutations for domestication traits appearing at different parts of the species range. The geographic structure in this domesticated species suggests that admixture might have allowed local domestication alleles to spread into other proto-domesticated *O. glaberrima* genetic groups in different parts of West and Central Africa, and facilitated the development of modern domesticated crop species which contain multiple domestication alleles sourced from different areas. Thus, in the end these gradual changes occurring across multiple regions provided different mutations at key domestication genes, which ultimately spread and came together to form modern *O. glaberrima*.

There has been intense debate on the nature of domestication, and recently (with particular emphasis on early Fertile Crescent domestication) discussion on whether this process proceeds in localized (centric) vs. a diffuse manner across a wider geographic area (non-centric) [75,84,86]. As we begin to use more population genomic data and whole genome sequences, as well as identify causal mutations associated with key domestication traits, we can begin to study the interplay between geography, population structure and the evolutionary history of specific domestication genes, and reconstruct the evolutionary processes that led to the origin and domestication of crop species. Moreover, a functional phylogeographic approach, as demonstrated here, could provide geographic insights into key traits that underlie species characteristics, and may allow us to understand how functional traits originate and spread across a landscape.

## Materials and Method

### Sample genome sequencing

*O. glaberrima* and *O. barthii* samples were ordered from the International Rice Research Institute and their accession numbers can be found in S1 Table. DNA was extracted from a seedling stage leaf using the Qiagen DNeasy Plant Mini Kit. Extracted DNA from each sample was prepared for Illumina genome sequencing using the Illumina Nextera DNA Library Preparation Kit. Sequencing was done on the Illumina HiSeq 2500 – HighOutput Mode v3 with 2×100 bp read configuration, at the New York University Genomics Core Facility. Raw FASTQ reads are available from NCBI biproject ID PRJNA453903.

### Reference genome based read alignment

Raw FASTQ reads from the study Wang et al. [12] and Meyer et al. [13] were downloaded from the sequence read archive (SRA) website with identifiers SRP037996 and SRP071857 respectively.

FASTQ reads were preprocessed using BBTools (https://jgi.doe.gov/data-and-tools/bbtools/) bbduk program version 37.66 for read quality control and adapter trimming. For bbduk we used the option: minlen=25 qtrim=rl trimq=10 ktrim=r k=25 mink=11 hdist=1 tpe tbo; which trimmed reads below a phred score of 10 on both sides of the reads to a minimum length of 25 bps, 3’ adapter trimming using a kmer size 25 and using a kmer size of 11 for ends of read, allowing one hamming distance mismatch, trim adapters based on overlapping regions of the paired end reads, and trim reads to equal lengths if one of them was adapter trimmed.

FASTQ reads were aligned to the reference *O. glaberrima* genome downloaded from EnsemblPlants release 36 (ftp://ftp.ensemblgenomes.org/pub/plants/). Read alignment was done using the program bwa-mem version 0.7.16a-r1181 [87]. Only the 12 pseudomolecules were used as the reference genome and the unassembled scaffolds were not used. PCR duplicates during the library preparation step were determined computationally and removed using the program picard version 2.9.0 (http://broadinstitute.github.io/picard/).

### Sequence alignment analysis

Using the BAM files generated from the previous step, genome-wide read depth for each sample was determined using GATK version 3.8–0 (https://software.broadinstitute.org/gatk/).

Because of the differing genome coverage between samples generated from different studies, depending on the population genetic method we used different approaches to analyze the polymorphic sites. A complete probabilistic framework without hard-calling genotypes, was implemented to analyze levels of polymorphism (including estimating θ,Tajima’s D, and F,_ST_), population relationships (ancestry proportion estimation and phylogenetic relationship), and admixture testing (ABBA-BABA test). For methods that require genotype calls, we analyzed samples that had greater then 10× genome coverage. Details are shown below.

### Polymorphism analysis

We used ANGSD version 0.913 [50] and ngsTools [51] which uses genotype likelihoods to analyze the polymorphic sites in a probabilistic framework. ngsTools uses the site frequency spectrum as a prior to calculate allele frequencies per site. To polarize the variants the *O. rufipogon* genome sequence [33] was used. Using the *O. glaberrima* genome as the reference, the *O. rufipogon* genome was aligned using a procedure detailed in Choi et al. [34]. For every *O. glaberrima* genome sequence position, the aligned *O. rufipogon* genome sequence was checked, and changed to the *O. rufipogon* sequence to create an *O. rufipogon*-ized *O. glaberrima* genome. Gaps, missing sequence, and repetitive DNA were noted as ‘N’. For all analysis we required the minimum base and mapping quality score per site to be 30. We excluded repetitive regions in the reference genome from being analyzed, as read mapping to these regions can be ambiguous and leading to false genotypes.

The site frequency spectrum was then estimated using ANGSD with the command:

~~~
angsd -b $BAM_list -ref $Reference_Genome -anc $Outgroup_genome \
 -out $SFS \
 -uniqueOnly 1 -remove_bads 1 -only_proper_pairs 1 -trim 0 -C 50 -baq 1 \
 -minMapQ 30 -minQ 30 \
 -minInd $minInd -setMinDepth $setMinDepth -setMaxDepth $setMaxDepth \
 -doCounts 1 -GL 1 -doSaf 1
~~~

For each genetic group a separate site frequency spectrum was estimated and the options –minInd, –setMinDepth, and –setMaxDepth were changed accordingly. Parameter minInd represent the minimum number of individuals per site to be analyzed, setMinDepth represent minimum total sequencing depth per site to be analyzed, and setMaxDepth represent maximum total sequencing depth per site to be analyzed. We required –minInd to be 80% of the sample size, –setMinDepth to be one-third the average genome-wide coverage, and –setMaxDepth to be 2.5 times the average genome-wide coverage. Using the site frequency spectrum, θ was calculated with the command:

~~~
angsd -b $BAM_list -ref $Reference_Genome -anc $Outgroup_genome \
 -out $Theta \
 -uniqueOnly 1 -remove_bads 1 -only_proper_pairs 1 -trim 0 -C 50 -baq 1 \
 -minMapQ 30 -minQ 30 \
 -minInd $minInd -setMinDepth $setMinDepth -setMaxDepth $setMaxDepth \
 -doCounts 1 -GL 1 -doSaf 1 -doThetas 1 -pest $SFS
~~~

The θ estimates from the previous command was used to compute sliding window values for Tajima’s θ and D [63] with the command:

~~~
thetaStat do_stat $Theta -nChr $Indv -win 10000 -step 10000
~~~

The option nChr is used for the total number of samples in the group being analyzed. Window size was set as 10,000 bp and was incremented in non-overlapping 10,000 bp.

F_ST_ values between pairs of population were also calculated using a probabilistic framework. Initially, we calculated the joint site frequency spectrum (2D-SFS) between the two populations of interest with the command:

~~~
realSFS $Pop1_SFS $Pop2_SFS > $Pop1_Pop2_2DSFS
~~~

Each population’s site frequency spectrum estimated from previous step is used to estimate the 2D-SFS. With the 2D-SFS F_ST_ values were calculated with the command:

~~~
realSFS fst index $Pop1_SFS $Pop2_SFS -sfs $Pop1_Pop2_2DSFS \
 -fstout $Pop1_Pop2_Fst
~~~

F_ST_ values were calculated in non-overlapping 10,000 bp sliding windows. For the sliding windows calculated for θ, Tajima’s D, and F_ST_ values, we required each window to have at least 30% of the sites with data or else the window was discarded from being analyzed.

### ABBA-BABA test

Evidence of admixture between genetic groups were determined with a multi-population ABBA-BABA test [60], using the abbababa2 function in ANGSD:

~~~
angsd -doAbbababa2 1 -bam “$BAMLIST” -sizeFile $SizeFile \
 -out $ABBA_BABA_out \
 -anc $Outgroup_genome -maxDepth 6000 -doCounts 1 \
 -useLast 0 -minQ $minQ -minMapQ $minMapQ
~~~

The *O. rufipogon*-ized *O. glaberrima* genome was used as the outgroup genome sequence. Because the topology of the relationship between OG-B and OG-C was uncertain, we conducted ABBA-BABA tests assuming OG-A and OG-B was sister groups, or assuming OG-A and OG-C was sister groups. The allele counts from the abbababa2 function were analyzed with the estAvgError.R R script [88], which was supplied from the ANGSD package, to calculate D-statistics and the Z-score. The Z-scores were calculated from using a block jack-knife approach where the whole genome divided into 5 Mbp blocks.

### Determining population relationships

Ancestry proportions were estimated using NGSadmix [52]. Initially, genotype likelihoods were calculated using ANGSD with the command:

~~~
angsd -b $BAM_list -ref $Reference_Genome -anc $Outgroup_genome \
 -out $GL \
 -uniqueOnly 1 -remove_bads 1 -only_proper_pairs 1 -trim 0 -C 50 -baq 1 \
 -minMapQ 30 -minQ 30 \
 -minInd $minInd -setMinDepth $setMinDepth -setMaxDepth $setMaxDepth \
 doCounts 1 -GL 1 -doMajorMinor 4 -doMaf 1 \
 -skipTriallelic 1 -doGlf 2 -SNP_pval 1e-3
~~~

To reduce the impact of LD would have on the ancestry proportion estimation, we randomly picked a polymorphic site in non-overlapping 50 kbp windows. In addition we made sure that the distance between polymorphic sites were at least 25 kbp apart. We then used NGSadmix to estimate the ancestry proportions for K=2 to 9. For each K the analysis was repeated 100 times and the ancestry proportion with the highest log-likelihood was selected to represent that K.

Phylogenetic relationships between samples were reconstructed using the genetic distance between individuals. Distances were estimated using genotype posterior probabilities from ANGSD command:

~~~
angsd -b $BAM_list -ref $Reference_Genome -anc $Outgroup_genome \
 -out $GPP \
 -uniqueOnly 1 -remove_bads 1 -only_proper_pairs 1 -trim 0 -C 50 -baq 1 \
 -minMapQ 30 -minQ 30 \
 -minInd $minInd -setMinDepth $setMinDepth -setMaxDepth $setMaxDepth \
 doCounts 1 -GL 1 -doMajorMinor 4 -doMaf 1 \
 -SNP_pval 1e-3 -doGeno 8 -doPost 1
~~~

Genotype posterior probability was used by NGSdist [55] to estimate genetic distances between individuals, which was then used by FastME ver. 2.1.5 [89] to reconstruct a neighbor-joining tree. Tree was visualized using the website iTOL ver. 3.4.3 (http://itol.embl.de/) [90].

### SNP calling

Since several methods require genotype calls for analysis SNP calling was also performed. Samples with greater then or equal to 10× genome coverage (GE10 dataset) was considered to ensure sufficient read coverage for each site at the cost of excluding individuals from genotype calling. These were 174 individuals that belonged to the genetic grouping designated by this study, and full list of individuals can be found in S9 Table.

For each sample, genotype calls for each site was conducted using the GATK HaplotypeCaller engine under the option `-ERC GVCF` mode to output as the genomic variant call format (gVCF). The gVCFs from each sample were merged together to conduct a multi-sample joint genotyping using the GATK GenotypeGVCFs engine. Genotypes were divided into SNP or INDEL variants and filtered using the GATK bestpractice hard filter pipeline [91]. For SNP variants we excluded regions that overlapped repetitive regions and variants that were within 5 bps of an INDEL variant. We then used vcftools version 0.1.15 [92] to select SNPs that had at least 80% of the sites with a genotype call, and exclude SNPs with minor allele frequency <2% to remove potential false positive SNP calls arising from sequencing errors or false genotype calls.

### Treemix analysis

Population relationships were examined as admixture graphs using the method in Treemix version 1.13 [57]. SNP calls from the GE10 dataset was used to calculate the allele frequencies for each genetic group. One thousand SNPs were analyzed together as a block to account for the effects of LD between SNPs. The OB-W genetic group was used as the outgroup and a Treemix model assuming 0–3 migration events were fitted. The four-population test [58] was conducted using the fourpop program from the Treemix package.

### Estimating levels of linkage disequilibrium

Genome-wide levels of LD (r^2^) was estimated with the GE10 dataset and using the program PLINK version 1.9 [93]. LD was calculated for each genetic group separately across a non-overlapping 1Mbp window and between variants that are at most 99,999 SNPs apart. LD data was summarized by calculating the mean LD between a pair of SNPs in 1,000 bp bins. A LOESS curve fitting was applied for a line of best fit and to visualize the LD decay.

### Past demography estimation

Past effective population size changes were estimated using the method in PSMC’ (MSMC model for two haplotypes) [64]. BAM files for the samples from the GE10 dataset was used to call genotypes with the mpileup command in samtools version 1.3.1. Sites with a minimum base and mapping quality scores of 30 were analyzed while genotypes from repetitive regions were excluded. The generate_multihetsep.py script from the msmc-tools suite (https://github.com/stschiff/msmc-tools) was used to convert the VCF file into the input format for MSMC. Due to the inbreeding, each individual was considered as a single genomic haplotype and individuals within the same genetic group were paired up to form a pseudodiploid [67]. All pairwise pseudodiploid combinations within the same genetic group was created to examine the coalescence within the same genetic groups, and each pseudodiploid PSMC’ profile was considered a biological replicate. The mutation scaled time and effective population sizes from the PSMC’ output were converted assuming a mutation rate of 6.5 × 10^-9^ substitutions per site per year [94] and a generation time of one generation per year.

### Determining gene orthologs between Asian and African Rice

We downloaded protein coding sequences for *O. sativa, O. glaberrima*, and *O. barthii* from EnsemblPlants release 36. An all-vs-all reciprocal BLAST hit approach was used to determine orthologs between species and paralogs within species. We used the program Orthofinder ver. 1.19 [95] to compare the proteomes between and within species for ortholog assignment. Orthofinder used the program DIAMOND ver. 0.8.37 [96] for sequence comparisons.

### Gene deletion analysis of genes *sh1* and *PROG1*

Synteny based on the *O. sativa sh1* gene (*Ossh1*; *O. sativa cv. japonica* chromosome 3:25197057–25206948) indicated orthologs surrounding *Ossh1* was found in chromosome 3 of *O. barthii* and on an unassembled scaffold named Oglab03_unplaced035 in *O. glaberrima* (S10 Table). The *sh1* gene was missing in *O. glaberrima* suggesting the gene deletion may have led to complex rearrangements that prevented correct assembly of the region in the final genome assembly. Because of this we used the *O. barthii* genome sequence to align raw reads and call polymorphic sites for downstream analysis.

The approximate region of the deletion in the *O. barthii* genome coordinate was examined by looking at the polymorphic sites, since our quality control filter removed polymorphic sites if it had less than 80% of the individuals with a genotype call. Between the genomic positions at *O. barthii* chromosome 23,100,000–23,130,000, no polymorphic sites passed the quality control filter (S13 Fig) and contained the gene *Obsh1*. Between the region at *O. barthii* chromosome 7:2,655,000–2,675,000 there was also no polymorphisms passing the filter and contained the gene *ObPROG1* (S14 Fig).

Gene deletion was inferred from comparing the read depth of a genic region inside and outside a candidate deletion region. Read depth was measured using bedtools ver. 2.25.0 [97] genomecov program. Individuals with and without the deletion were determined by comparing the median read coverage of the domestication gene within the candidate deletion region, to a gene that is outside the deletion region. We checked the orthologs to make sure the gene outside the deletion region existed in *O. barthii, O. glaberrima*, and *O. sativa*. To determine the *sh1* deletion status we examined its read depth and compared it to the *O. barthii* gene OBART03G27620 that was upstream and outside the candidate deletion region. Ortholog of OBART03G27620 is found in both *O. sativa* (Os03g0648500) and *O. glaberrima* (ORGLA03G0257300). To determine the deletion status of *PROG1* gene we examined its read depth and compared to *O. barthii* gene OBART07G03440. Ortholog of OBART07G03440 is found in both *O. sativa* (Os07g0153400) and *O. glaberrima* (ORGLA07G0029300).

Because some individuals had low genome-wide coverage (S1 Table) there is the possibility that some of those individuals had been detected as false positive deletion events. There are two main reasons we believe the deletions are likely to be present even for low coverage individuals. For example for the *sh1* deletion, (i) all individuals had at least a median coverage of ∼1× in the OBART03G27620 gene (S11 Table) suggesting read coverage may be low but if the gene is not deleted it is evenly distributed across a gene, and (ii) even comparing individuals with and without the *sh1* deletion that had a ∼1× median coverage in the non-deleted OBART03G27620 gene, there were clear differences in the *sh1* gene coverage (S15 Fig) where the individuals with the deletion always had a median coverage of zero.

### Shattering gene nomenclature

Gene names for the non-shattering phenotype have unfortunately varied between different *Oryza* studies. Genetic studies comparing Asian rice *O. sativa cv*. Japonica and its wild progenitor *O. rufipogon* had identified a single dominant allele responsible for non-shattering and named the locus as *Sh3* [98,99]. The causal gene was later identified on chromosome 4 and was given a new name as *sh4* [14]. Studies have used the names *Sh3* and *sh4* synonymously as the common gene name for the gene with locus ID Os04g0670900 [72].

Lu et al. had found an *O. glaberrima* specific gene deletion in chromosome 3 that caused a non-shattering phenotype and named this gene as *SH3. SH3* belongs to a YABBY protein family transcription factor. Using the *SH3* coding sequence in *O. barthii* (*ObSH3*), which the gene is not deleted, orthologs were found in maize (B4FY22), barley (M0YM09), and Brachypodium (I1GPY5) (Lu et al. In press). We discovered this group of proteins belonged to a group identified in Plant Transcription Factor Database ver 4.0 [100] under the ID OGMP1394. The *O. sativa* gene member of this group was gene ID Os03g0650000, which has previously been identified as a gene involved in non- shattering [101]. Thus, *ObSH3* and Os03g0650000 are orthologs of each other and Os03g0650000 has been named as *sh1*. Here, we followed the guideline recommended by Committee on Gene Symbolization Nomenclature and Linkage (CGSNL) [102] to designate *SH3* from Lu et al. as *sh1* to avoid using the overlapping gene name *sh3*.

### Gene haplotype analysis

To investigate the haplotype structure around the domestication genes we used all individuals from *O. glaberrima*, OB-G, and OB-W population regardless of the genome coverage. The *O. glaberrima* and *O. barthii* genome were used as reference to align the raw reads and call polymorphisms as outlined above. Missing genotypes were then imputed using Beagle version 4.1 [103].

We used vcftools to extract polymorphic sites around a region of interest. The region was checked for evidence of recombination using a four-gamete test [74], to limit the edges connecting haplotypes as mutation distances during the haplotype network reconstruction. To minimize false positive four-gamete test results caused from technical errors such as genotype error and sequencing error, if the observed frequency of the fourth haplotype was below 1% we considered the haplotype an error and did not consider it as evidence of recombination. If a region had evidence of recombination we checked if the recombination was limited to the wild or domesticated African rice. If recombination was only detected in the wild population then we determined the pair of SNPs that failed the four-gamete test. Here, because the four-gamete test did not detect any evidence of recombination in the *O. glaberrima* population, the fourth haplotype observed in the wild population is only limited to *O. barthii* and do not provide any information with regard to the direct origin of the *O. glaberrima* haplotypes. Hence, we removed individuals with the fourth haplotype and estimated the haplotype network of the region.

Haplotype network was reconstructed using the R pegas [104] and VcfR [105] package, using the hamming distance between haplotypes to construct a minimum spanning tree. We also reconstructed a neighbor-joining tree of the same region that used to build the haplotype network, as an independent method to validate the results. Pairwise distances between individuals were measured using the Kronecker delta function [106]. Using the distance matrix neighbor-joining tree was built using FastME and plotted with iTOL.

## Acknowledgements

We thank the New York University Genomics Core Facility for sequencing support and the New York University High Performance Computing for supplying the computational resources.

## Supporting information

**S1 Fig. Geographic distribution of analyzed *O. glaberrima* and *O. barthii* samples.** (A) *O. glaberrima* from this study. (B) *O. glaberrima* from Meyer et al. (C) *O. barthii* samples. Note for the majority of *O. barthii* samples from Wang et al. the locations are unknown.

**S2 Fig. Ancestry proportion estimates for K=2 to 9.** Black stars below the admixture barplot indicate *O. glaberrima* individuals. Colored stars above admixture barplot are the *O. barthii* grouping designated by Wang et al. where blue: OB-I, brown: OB-II, red: OBIII, yellow: OB-IV, and pink: OB-V group.

**S3 Fig. Neighbor-joining tree built using a distance matrix estimated from NGSdist.** Color strips represent the *O. barthii* grouping designated by Wang et al.

**S4 Fig. Ancestry proportion estimates for K=2, 5, and 7.** Black stars below the admixture barplot indicate the two *O. glaberrima* individuals IRGC103631 and IRGC103638. Colored stars above admixture barplot are the *O. barthii* grouping designated by Wang et al. where blue: OB-I, brown: OB-II, red: OB-III, yellow: OB-IV, and pink: OB-V group

**S5 Fig. Treemix admixture graphs for (A) no migration, (B) 1 migration, and (C) 2 migration models.** Residuals between genetic groups are shown under the admixture graph.

**S6 Fig. Levels of polymorphism and Tajima’s D for OB-G, OB-W and *O. glaberrima*.** Significant difference after Mann-Whitney U test (p < 0.001) are indicated with three stars.

**S7 Fig. PSMC’ estimated demography changes in OB-W genetic group from pseudo-diploids generated from OB-I and OB-II group.**

**S8 Fig. Mid-point rooted neighbor-joining tree of the *sh4* region in Fig 4A.** Star indicates the individuals without the nonsense mutation. Individuals from haplogroup I from Fig 4B are shown in grey boxes.

**S9 Fig. Mid-point rooted neighbor-joining tree of the upstream and downstream of *sh1* deletion region.**

**S10 Fig. π_w_/ π_D_ statistics around the *PROG1* region in *O. barthii* and *O. glaberrima* reference genomes.**

**S11 Fig. Mid-point rooted neighbor-joining tree of the upstream and downstream 5kbp of *PROG1* deletion region.** Individuals from haplogroup IV from Fig 4F are shown in grey boxes.

**S12 Fig. Haplotype network of the downstream 5 kbp of the *PROG1* deletion.**

**S13 Fig. Genome coordinate of chromosome 3 and presence of a polymorphism is indicated with a point.**

**S14 Fig. Genome coordinate of chromosome 7 and presence of a polymorphism is indicated with a point.**

**S15 Fig. Visualization of read pileup of a region upstream and a region within the *Obsh1* gene for individuals with and without the *sh1* deletion.** In each panel an individual with low and high coverage are compared.

**S1 Table. Information on the sequenced and analyzed individuals of this study.**

**S2 Table. Count of country of origin for individuals from OB-G and OB-W genetic group.**

**S3 Table. ABBA-BABA test result testing admixture between OB-W and *O. glaberrima* genetic groups.**

**S4 Table. Individuals corresponding to the haplotype groups identified in Fig 4A *sh4* gene region.**

**S5 Table. Individuals corresponding to the haplotype groups identified in Fig 4B *sh1* gene region.**

**S6 Table. *O. glaberrima* and *O. barthii* genes syntenic to the *O. sativa PROG1* region.**

**S7 Table. Individuals corresponding to the haplotype groups identified in Fig 4C *PROG1* upstream gene region.**

**S8 Table. Individuals corresponding to the haplotype groups identified in S12 Fig *PROG1* downstream gene region.**

**S9 Table. Individuals with greater then 10× genome coverage.**

**S10 Table. *O. glaberrima* and *O. barthii* genes syntenic to the *O. sativa sh1* region.**

**S11 Table. Read coverage count for *sh1* and *PROG1* region.**

